# Multiple glycoforms of TrkA interact with N-cadherin during trigeminal ganglion neurodevelopment

**DOI:** 10.1101/2025.11.14.688512

**Authors:** Caroline A. Halmi, Lisa A. Taneyhill

## Abstract

The trigeminal ganglion is a component of the sensory nervous system that arises from neural crest and placode cells. The dual origin of the trigeminal ganglion leads to a heterogenous neuronal population that transmits somatosensory information from the face back to the brain. Proper trigeminal ganglion development relies, in part, on neurotrophic signaling, including interactions between Nerve Growth Factor and its cognate receptor, Tropomyosin receptor kinase A (TrkA), a receptor tyrosine kinase. Post-translational modifications, including glycosylation, play a crucial role in the ability of TrkA to reach the plasma membrane, yet the specific glycan profile and functional relevance of these modifications have not been characterized in sensory neurons *in vivo*. Here, we sought to characterize the different glycosylation events occurring on TrkA during trigeminal ganglion neurodevelopment. We discovered that multiple glycoforms of TrkA exist that correlate to partially and fully mature versions of the protein reported *in vitro*. Furthermore, we discovered that TrkA interacts with a cell adhesion molecule, N-cadherin, on membranes of trigeminal neurons, both in the cell bodies and axons. Based on the size of the TrkA bands that interact with N-cadherin, our results suggest these interactions are occurring both on the plasma membrane and intracellularly on the membrane of organelles. While interactions between receptor tyrosine kinases and cadherins have been shown in other contexts, our findings are the first to identify such an interaction in the trigeminal ganglion and suggest an important role for coordination between neurotrophic signaling and cell adhesion for proper neurodevelopment, both during TrkA protein maturation and during receptor tyrosine kinase signaling. Given that aberrant receptor tyrosine kinase and cadherin signaling is commonly implicated in neurodevelopmental disorders and cancer, understanding how these interactions are established during normal development may provide additional insight into their dysregulation during disease.

## Introduction

Cranial ganglia are components of the peripheral nervous system that transmit sensory information from the surrounding environment back to the central nervous system (Schlosser, 2018). The trigeminal ganglion, the largest of all cranial ganglia, extensively innervates the face to relay somatosensory information including pain, touch, and temperature (Heimer, 1983). It is derived from two embryonic cell populations, neural crest cells and placode cells, which originate in different anatomical locations but eventually become migratory and coalesce to form this sensory structure (D’Amico-Martel & Noden, 1980; Hamburger, 1961). While both cell types differentiate into neurons, neural crest cells also give rise to supporting glia. This dual embryonic origin of trigeminal sensory neurons leads to a heterogenous population with distinct molecular profiles and signaling responses (Bhuiyan et al., 2024; Marmigère & Ernfors, 2007; M. Q. Nguyen et al., 2017).

Among the signaling pathways that are critical for proper trigeminal ganglion development are those initiated downstream of neurotrophins and their cognate receptors, which regulate neuronal survival, outgrowth, and subtype specification (Barbacid, 1995; Cohen et al., 1954; Levi-Montalcini & Hamburger, 1953). As development progresses and trigeminal neurons respond to environmental cues, specific subclasses of neurons emerge including nociceptors, mechanoreceptors, and thermoreceptors that detect pain, mechanical stimuli, and temperature, respectively. These neuronal subtypes can be identified in part by the expression of members of the receptor tyrosine kinase (RTK) family known as the tropomyosin receptor kinases (Trks), which are activated by secreted neurotrophic factors (Davies, 1997; Ernfors et al., 1992; Huang, Wilkinson, et al., 1999; Huang, Zang, et al., 1999; Mu et al., 1993).

TrkA, the high-affinity receptor for nerve growth factor (NGF), is expressed in the majority of nociceptive neurons (Fang et al., 2005). Upon ligand binding and subsequent receptor dimerization, TrkA undergoes autophosphorylation to activate downstream signaling pathways like MAPK, PI3K/Akt, and PLCγ, which promote neuronal survival and differentiation (Huang & Reichardt, 2003; Marlin & Li, 2015). Loss of NGF-TrkA signaling leads to increased apoptosis in small diameter neurons of both the trigeminal ganglion and dorsal root ganglion, confirming its essential role in neuronal survival and health during neurodevelopment (Johnson et al., 1980; Ruit et al., 1992; Smeyne et al., 1994). Although the function of TrkA has been characterized extensively in rat pheochromocytoma cells (PC12) (Negrini et al., 2013; Zhou et al., 1995) and models such as rodent (Fang et al., 2005; McMahon et al., 1994) and chick dorsal root ganglia (Gallo et al., 1997; George et al., 2010; Karlsson et al., 1998; Rifkin et al., 2000; Williams et al., 1995), far less is known about the post-translational mechanisms that regulate TrkA maturation, trafficking, and localization to the cell surface, which are critical for its function as a receptor. Existing information on these processes has been obtained exclusively from studies in PC12 cells, leaving these mechanisms unexplored *in vivo* within cranial sensory ganglia.

Proper localization of TrkA to the plasma membrane relies on a series of regulated processing and maturation steps (Jullien et al., 2002; Martin-Zanca et al., 1989; Watson et al., 1999; Zhou et al., 1995). Data from prior studies indicate that TrkA is synthesized as an immature, unmodified precursor protein. In PC12 cells, the unglycosylated core TrkA protein is approximately 75 kD, although whether this remains true *in vivo* is unknown (Watson et al., 1999). Next, TrkA undergoes glycosylation in the ER to become a 110 kD precursor glycoprotein that undergoes further glycosylation in the Golgi to yield a 140 kD mature form prior to its transit to the cell membrane. Previous studies have shown that the 110 kD, partially glycosylated form of TrkA is N-linked glycosylated, while the 140 kD form contains additional sialic residues, likely on these N-linked glycans. Recently, a report has also suggested the presence of O-linked glycosylation on TrkA in cancer cells, though its presence and functional significance, particularly during normal development, are unclear (Lin et al., 2022).

In addition to RTKs, cell adhesion molecules such as cadherins are critical for neuronal organization and signaling. Cadherins regulate cell-cell adhesion in a calcium-dependent manner and contribute to tissue morphogenesis by influencing proliferation and intracellular signaling (Halbleib & Nelson, 2006). Although multiple cadherins are noted in different tissues, neural cadherin (N-cadherin) is broadly expressed in the developing nervous system, including in trigeminal placode cells prior to their delamination, in their neuronal derivatives, and later in all trigeminal sensory neurons irrespective of cellular origin (Hatta & Takeichi, 1986; Shiau et al., 2008). Additionally, N-cadherin knockdown in trigeminal placode cells disrupts initial trigeminal ganglion condensation and later axon outgrowth through cell- and non-cell autonomous mechanisms involving adhesion (Halmi et al., 2025; Shiau & Bronner-Fraser, 2009). Beyond their adhesive roles, cadherins can also interact with signaling receptors, including RTKs, to functionally regulate downstream signaling, receptor membrane localization, and cell behaviors such as migration and proliferation (Andl & Rustgi, 2005; Madarampalli et al., 2019; T. Nguyen & Mège, 2016; Qian et al., 2004). Dysregulation of these RTK-cadherin interactions are commonly seen in diseases such as cancer, where they contribute to the unchecked proliferation of cells. However, whether such RTK-cadherin interactions occur normally in the developing chick trigeminal ganglion, and how these interactions might influence trafficking and localization of receptors, has not yet been explored.

In this study, we investigate the glycosylation status of TrkA in chick trigeminal sensory neurons, including the relationship between TrkA and N-cadherin. Our results demonstrate that distinct TrkA glycoforms interact with N-cadherin in cell bodies versus axons, and these interactions occur on neuronal cellular membranes, likely on the plasma membrane and the membranes of organelles. Our findings reveal a previously uncharacterized interaction between neurotrophic signaling factor receptors and cell adhesion molecules that may regulate normal trafficking of TrkA to permit proper neuronal differentiation and survival during sensory ganglion development.

## Methods

### Chick embryos

Fertilized chicken eggs (*Gallus gallus*) were obtained from the Department of Animal and Avian Sciences at the University of Maryland (MD) and incubated at 37°C in humidified incubators on their sides. Embryos were incubated to reach desired Hamburger Hamilton (HH) stages based on times provided therein (Hamburger & Hamilton, 1992).

### Immunohistochemistry

Embryos were collected at desired stages of interest and fixed in 4% paraformaldehyde (PFA) overnight at 4°C with agitation. After fixation, embryos were embedded and sectioned as described previously (Halmi et al., 2025). Briefly, embryos were processed through increasing concentrations of sucrose (5% and 15%) and gelatin (7.5% and 20%), before being embedded in 20% gelatin. Embedded embryos were cryo-sectioned and immunostaining was performed (Halmi et al., 2025). Primary antibodies used were the following: anti-TrkA (a gift from Dr Francis Lefcort, Montana State University, Bozeman, MT, USA; 1:5000) and anti-N-cadherin (6B3, DSHB, 1:75). Secondary antibodies (goat anti-rabbit IgG and goat anti-mouse IgG_1_) were used at 1:250.

### Trigeminal ganglion explant cultures

Wild-type embryonic day (E)6.5/HH28-30 embryos were removed from eggs and stored on ice in neuronal dissection media (1X Hank’s Balanced Salt solution with 1% pen/strep, 10 mM HEPES, and 1.4 mM glucose). Trigeminal ganglia were dissected on a sylgard-coated dish by first bisecting the embryo and then removing the trigeminal ganglion and axonal projections with tungsten needles. Dissected ganglia were dispersed with a P10 pipette and cultured on two-well glass chamber slides (ThermoFisher) coated with Poly-L-Lysine (0.01% PLL, Sigma) and fibronectin (0.01%, Sigma). To make the coated slides, PLL was added to each chamber on the glass slides and incubated inside a biosafety cabinet for 30 minutes at room temperature. The PLL was removed, and slides were washed three times with water. Next, fibronectin diluted in DMEM media (ThermoFisher) was added to each chamber and incubated at 37°C for a minimum of 2 hours. The fibronectin solution was removed, and slides were stored at 4°C until needed (for up to 1 week). Explants were grown in Neurobasal complete media (ThermoFisher) supplemented with B27 (1%, ThermoFisher), N2 (0.5%, ThermoFisher), Glutamax (1%, ThermoFisher), penicillin/ streptomycin (1%, ThermoFisher), and with nerve growth factor (ThermoFisher), brain-derived neurotrophic factor (Amgen) and neurotrophin-3 (Amgen), all at 50 ng/ml.

### Explant immunocytochemistry

Explant slides were washed once with 1X PBS, followed by fixation in 4% PFA for 30 minutes at room temperature with agitation. After fixation, slides were rinsed with 1X PBS and stored in 1X PBS at 4°C until ready for immunocytochemistry. A hydrophobic perimeter was drawn around the slides using an ImmEdge Pen (Vector Laboratories) to prevent dehydration of cells, and explants were taken through blocking, primary antibody incubation, and secondary antibody incubation in the absence of coverslips, as described in the immunohistochemistry methods. Slides were mounted with DAPI-fluoromount media (SouthernBiotech) and covered with a glass coverslip.

### Confocal imaging

All images were acquired with the LSM Zeiss 800 confocal microscope. Laser power, gain and offset were kept consistent for the different channels during all experiments where possible. Adobe Photoshop CC 2025 was used for image processing.

### Immunoprecipitation

Wild-type E6.5/HH28-30 trigeminal ganglia were dissected from embryos and cell bodies were separated from axons using fine tungsten needles. Samples were centrifuged for 5 minutes at 4°C at 2292 x*g*. Pellets were flash frozen in liquid nitrogen and stored at -80°C. When needed, samples were lysed in lysis buffer (50 mM Tris (pH 7.5), 100 mM NaCl and 0.5% IGEPAL CA-630)) supplemented with cOmplete Protease Inhibitor Cocktail (Roche) and 1 mM PMSF for 30 minutes on ice with periodic gentle vortexing. Lysates were then centrifuged at 20,627 x*g* for 20 minutes at 4°C. The soluble protein fraction was then quantified by Bradford assay (ThermoFisher). N-cadherin-TrkA coimmunoprecipitations were performed with Protein A/G magnetic beads (ThermoFisher) according to the manufacturer’s instructions. Briefly, equivalent amounts of protein lysates were incubated overnight with 10 µg of either the N-cadherin antibody (Abcam, ab182) or a rabbit IgG antibody (R&D Systems, AB-105-C) at 4°C with constant mixing. The following day, the lysates were combined with washed magnetic beads (3 times in wash buffer) and incubated for 1 hour at room temperature with constant mixing. After this, the bound proteins were washed on the beads three times before being eluted off by boiling at 99°C for 7 minutes in 1X reducing Laemmli sample buffer.

### Immunoblotting

Samples were separated by 8% SDS-PAGE, then transferred to a nitrocellulose membrane (BioRad). The membrane was blocked in 5% milk (in 1X PBS+0.1% Tween-20 (PTW)) for 30 minutes at room temperature followed by overnight incubation at 4°C with the following primary antibodies diluted in blocking solution: N-cadherin (MNCD2, DSHB 1:100) and TrkA (MSU, 1:5000 or Origene, 1:1000). Membranes were washed three times for 10 minutes each in PTW before incubation with secondary antibodies (species- and isotype-specific horseradish peroxidase, Jackson ImmunoResearch) diluted in blocking solution for 1 hour at room temperature. Membranes were washed three times for 10 minutes each in PTW. Antibody detection was performed using Supersignal West Pico or Femto chemiluminescent substrates (ThermoFisher) and visualized using a ChemiDoc XRS system (Bio-Rad).

To calculate protein molecular weights, a straight vertical line was drawn through the protein ladder and a profile plot was generated on Fiji (Schindelin et al., 2012). The pixel value for each peak, which corresponded to a molecular weight marker was recorded. The log_(10)_ value for each molecular weight marker was calculated and a graph was generated with the migration distance (pixel value) along the x-axis and the log_(10)_ value along the y-axis. A linear regression was fitted and the corresponding equation of the line was used to calculate the molecular weight of all unknown bands.

### Enzymatic treatments of lysate

For all enzymatic treatments, equal amounts of trigeminal ganglia lysate were used. PNGase F (NEB, P0704S), O-glycosidase (NEB, P0733S), and α2-3,6,8 Neuraminidase (NEB, P0720S) were added to lysate per manufacturer’s guidelines. Samples were incubated for 2-4 hours at 37°C. After incubation, 1X reducing Laemmli sample buffer was added to each sample, and samples were boiled at 99°C for 7 minutes before protein separation by SDS-PAGE.

### Glycoprotein enrichment

#### Concanavalin A (ConA) and Wheat Germ Agglutinin (WGA)

To enrich for N-linked glycans and sialic acid residues, ConA and WGA Glycoprotein Isolation kits were used, respectively (ThermoFisher, 89804, 89805). Briefly, the resin was first washed three times with 1X Wash Buffer. Trigeminal ganglia lysate was added to the washed resin and 1X Wash Buffer was added to reach a final volume of 250 µL. Lysate/resin mixtures were incubated for 1 hour at room temperature with constant mixing. After incubation, glycoprotein-bound resin was washed four times with 1X Wash Buffer. After the final wash, 1X reducing Laemmli sample buffer was added to each sample, and samples were boiled at 99°C for 7 minutes before protein separation by SDS-PAGE. This experiment was performed on unconjugated agarose resin as described above for a control.

#### Peanut Agglutinin (PNA)

To enrich for O-linked glycans, biotinylated PNA (Vector Laboratories, B-1075-5) was used. Prior to enrichment, the lysate was treated with α2-3,6,8 Neuraminidase (NEB, P0720S) to remove sialic acid residues. In a microcentrifuge tube, neuraminidase-treated lysate, biotinylated PNA (10 µg/mL), and Wash/Binding Buffer (1X PBS with 0.05% Tween-20) were combined to reach a volume of 500 µL and incubated overnight at 4°C with constant mixing. The next day, streptavidin Sepharose resin (ThermoFisher, 20349) was washed with Wash/Binding Buffer three times before adding the lysate/lectin solution to it. Lysate/resin mixtures were incubated for 1 hour at room temperature with constant mixing. After incubation, glycoprotein-bound resin was washed four times with Wash/Binding Buffer. After the final wash, 1X reducing Laemmli sample buffer was added to the sample, which was then boiled at 99°C for 7 minutes before protein separation by SDS-PAGE. This experiment was performed on Sepharose resin without biotinylated PNA as described above for a control.

### Inhibitor treatment of explant cultures

Wild-type E6.5/HH28-30 trigeminal ganglia explant cultures were grown for 1 day *in vitro* as described in the explant cultures section. Tunicamycin (Sigma, T7765), Brefeldin A (Sigma, B6542), and Monensin (Sigma, M5273) were resuspended in DMSO to obtain a concentration of 1 mg/mL. The next day, all inhibitors, and the DMSO control, were separately added to Neurobasal complete media with brain-derived neurotrophic factor (Amgen, 50 ng/ml) and neurotrophin-3 (Amgen, 50 ng/ml), at a final concentration of 1 µg/mL, and each inhibitor (or control) was added to the explant cultures for 4 hours. After incubation, the media was removed, and explants were washed in 1X PBS before either being fixed in 4% PFA for immunostaining or flash frozen in liquid nitrogen and stored at -80°C to be processed for immunoblotting.

### Extraction of membrane and cytosolic protein fractions from trigeminal ganglia

Membrane and cytosolic proteins from E6.5/HH28-30 trigeminal ganglia were isolated using the Mem-PER Plus Membrane Protein Extraction Kit (ThermoFisher 89842), per manufacturer’s guidelines. Briefly, flash-frozen trigeminal ganglia were thawed on ice and washed once with Cell Wash Solution. Permeabilization Buffer was added to trigeminal ganglia, mixed until a homogenous solution was obtained, and incubated for 10 minutes at 4°C with constant mixing. The solution was centrifuged at 16,000 x*g* for 15 minutes at 4°C to pellet permeabilized cells. The supernatant which contained the cytosolic protein fraction was removed and transferred to a new tube. Solubilization Buffer was then added to the pellet containing the membrane-bound proteins and mixed until a homogenous solution was obtained. The solution was incubated for 30 minutes at 4°C with constant mixing before being centrifuged at 16,000 x*g* for 15 minutes at 4°C. The supernatant containing membrane-associated proteins was transferred to a new tube before both fractions were quantified by Bradford assay for subsequent experimentation.

## Results

### TrkA possesses multiple glycoforms in the trigeminal ganglion

Previous work has demonstrated the glycosylation of TrkA in PC12 cells, which serve as an *in vitro* model of neuronal differentiation. However, TrkA glycosylation has not been studied *in vivo*. Thus, we first sought to establish what types of glycans were present on TrkA in chick trigeminal neurons. We treated trigeminal ganglia lysate with enzymes that remove specific sugar modifications on proteins (PNGase F: N-linked glycans, O-glycosidase: Core 1 and Core 3 O-linked glycans, and neuraminidase: sialic acids) and assessed changes to TrkA banding patterns by immunoblotting (Fig. 1A). In both the input and mock control lanes, the latter containing the enzymatic buffers added to the lysate without any enzymes, we observed six bands: the common doublet observed for TrkA, which we calculated to be 137 and 107 kD, as well as multiple lower molecular weight bands that were 93 kD, 76 kD, 59 kD, and 53 kD. Notably, in some experiments, we also observed a 223 kD TrkA band in the input that was reduced in molecular weight in the mock control treatment (Supplemental Fig. 1).

**Figure 1:**
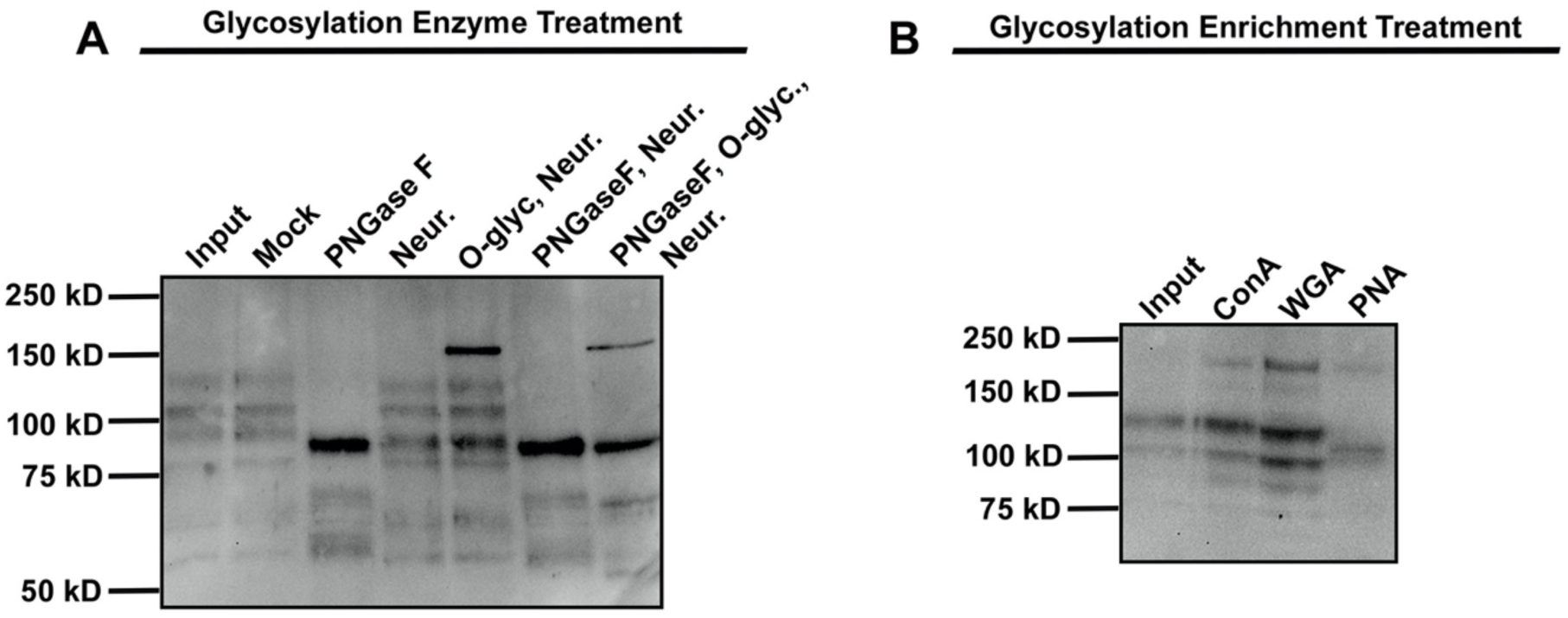
Different post-translational modifications occur on TrkA. (A) E6.5/HH28-30 trigeminal ganglia lysate treated with enzymes to inhibit glycosylation followed by immunoblotting for TrkA (N=3). Input: Untreated trigeminal ganglia lysate. Mock (control): Lysate mixed with enzyme buffers. PNGase F, Neuraminidase, O-glycosidase (O-glyc.). (B) E6.5/HH28-30 trigeminal ganglia lysate after glycan enrichment followed by immunoblotting for TrkA (N=3). Input: Untreated trigeminal ganglia lysate. Concanavalin A (ConA), Wheat Germ Agglutinin (WGA), Peanut Agglutinin (PNA).

Upon PNGase F treatment, we observed a total collapse of the 137 kD, 107 kD, and 93 kD bands to a singular 88 kD band, as well as two lower molecular weight bands at 67 kD and 56 kD, and the persistence of the 53 kD band. After neuraminidase treatment, we observed a slight shift in the 137 kD band down to 127 kD, and a slight reduction in size of the 93 kD band to 89 kD, but no change in the 107 kD, 76 kD, 59 kD, or 53 kD bands. Since sialic acid residues are often found at the end of O-glycans and mask the ability of O-glycosidase to cleave these glycans completely, O-glycosidase and neuraminidase treatment were performed together. Interestingly, we observed a prominent 177 kD band, in addition to the other band shifts observed in the neuraminidase only-treated lysate. When PNGase F and neuraminidase were used together, we observed the same results as the PNGase F only treatment. Lastly, in the combination treatment with PNGase F, O-glycosidase, and neuraminidase, we observed the 177 kD band we had seen previously after O-glycosidase + neuraminidase treatment, a prominent band at 88 kD, which we observed after PNGase F treatment, and two lower bands that were 63 kD and 52 kD in size (Fig. 1A). We also examined N-cadherin, whose banding patterns only changed in the presence of PNGase F (Supplemental Fig. 2). This is in line with other reports indicating that N-cadherin is only modified with N-linked glycans (Guo et al., 2009; Langer et al., 2012). Altogether, these enzyme treatments confirm that chick TrkA possesses multiple different types of sugars, including N-linked and sialic acid modifications, and indicate that the 107 kD TrkA band contains N-linked glycans while the 137 kD band is both N-linked and sialylated. Our results also suggest that O-linked glycosylation might be evident due to the presence of a 177 kD band after O-glycosidase + neuraminidase treatment.

To further explore the glycosylation profile of TrkA, we next performed experiments to enrich for different types of glycosylations through the use of Concavalin A (ConA), which binds alpha-linked mannose or terminal glucose residues that are associated with N-linked glycosylation; Wheat Germ Agglutinin (WGA), which recognizes N-acetyl-glucosamine groups or sialic acid; and Peanut Agglutinin (PNA), which binds galactosyl-(β-1,3)-N-acetyl-galactosamine, a component of the O-glycosylation pathway. Interestingly, TrkA was enriched after all three treatments, further supporting the presence of O-linked glycans on TrkA (Fig. 1B). In addition to the 137 kD, 107 kD, 93 kD, and 76 kD TrkA bands observed in Fig. 1A, ConA and WGA bound glycosylations on two higher molecular weight TrkA bands at 223 kD and 177 kD. PNA also pulled down the 223 kD band but did not enrich for as many bands as the ConA or WGA treatment. Instead, PNA bound a 117 kD form of TrkA, along with the 107 kD TrkA glycoform seen in other treatments. These experiments confirm that TrkA contains N-linked glycans and sialic acid residues, in line with data from PC12 cells, but also reveal, for the first time, additional O-linked glycosylation in chick sensory neurons. These results suggest that while similar post-translational modifications occur on TrkA *in vitro* and *in vivo*, additional modifications are also present in chick trigeminal ganglion neurons.

### Inhibiting components of the secretory pathway impacts TrkA processing

To further explore the role of glycan processing of TrkA, we treated trigeminal neuronal explant cultures with validated inhibitors of protein processing and assessed TrkA localization and expression via immunostaining and immunoblotting. We hypothesized that each inhibitor would yield a unique change in TrkA expression and/or localization depending upon which portion of the secretory pathway was perturbed. Although we observed multiple TrkA bands, we chose to focus only on the partially and fully glycosylated forms for our analyses, as these are the most prevalent forms that move through the secretory pathway. In the DMSO control-treated explants, we observed TrkA perinuclearly as well as along the cell membrane (Fig. 2A-A’), and the typical doublet (107 kD and 137 kD) observed for TrkA that correspond to the partially and fully glycosylated forms (Fig. 2E). We also observed the 223 kD TrkA form in all of our conditions. Interestingly, in these experiments, TrkA ran slightly differently, with the partially and fully glycosylated forms appearing at 98 kD and 122 kD, respectively. Monensin is an antibiotic that inhibits the export of proteins out of the Golgi (Griffiths et al., 1983; Hammerschlag et al., 1982). This treatment yielded an accumulation of TrkA in a perinuclear location, possibly the Golgi given its subcellular distribution and the mechanism of action of Monensin (Fig. 2B-B’). While we observed the typical banding pattern for TrkA by immunoblotting, both the 98 kD and 122 kD forms were reduced, with the fully glycosylated form of TrkA being more greatly impacted than the partially glycosylated form (6.5-fold decrease vs. 1.5-fold decrease, Fig. 2E-G). Brefeldin A is a fungal metabolite that inhibits the anterograde transport of proteins from the ER to the Golgi, causing proteins to accumulate in the ER (Chardin & McCormick, 1999). After Brefeldin A treatment, we observed diffuse expression of TrkA throughout the cytoplasm of neuronal cell bodies (Fig. 2C-C’). This result correlated with the presence of a very prominent 98 kD TrkA band after immunoblotting (7-fold increase) and a reduction in the intensity of the 122 kD band (10-fold decrease, Fig. 2E-G). Lastly, Tunicamycin is an antibiotic that inhibits N-linked glycosylation of proteins in the ER, leading to an accumulation of unfolded proteins and cellular stress (Struck et al., 1978; Takatsuki et al., 1971). After Tunicamycin treatment, we observed a reduction in overall expression of TrkA in the neuronal cultures and their axonal protrusions (Fig. 2D-D’). This phenotype is in line with what we observed via immunoblotting, in that overall TrkA levels were decreased (2-fold decrease in the 122 kD band, 5-fold decrease in the 98 kD band, Fig. 2E-G). Taken together, these experiments indicate that blocking the secretory pathway at distinct points alters TrkA distribution and expression, correlating with the portion of the pathway that was perturbed.

**Figure 2:**
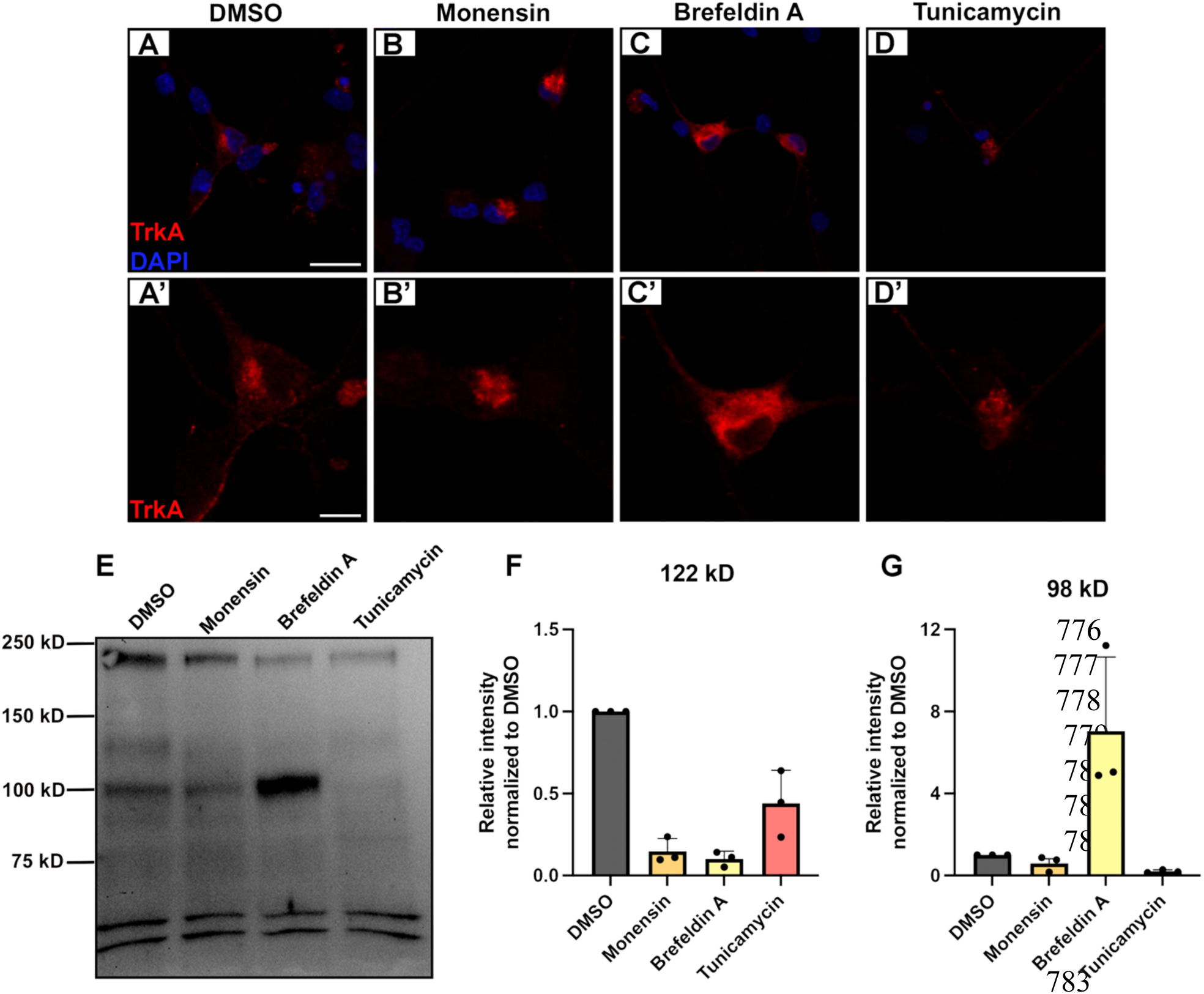
Inhibitors of the secretory pathway disrupt TrkA expression and glycosylation within trigeminal neurons. (A-D’) Representative images of cultured trigeminal ganglia following treatment with DMSO (A-A’), Monensin (B-B’), Brefeldin A (C-C’), or Tunicamycin (D-D’), and immunocytochemistry for TrkA and DAPI (marks nuclei). Scale bar in (A) is 25 µm and applies to (B, C, D). Scale bar in (A’) is 10 µm and applies to (B’, C’, D’). (E) Immunoblot for TrkA using lysate prepared from trigeminal ganglia explant cultures following treatment with DMSO, Monensin, Brefeldin A, and Tunicamycin. (F-G) Quantification of 122 kD (F) and 98 kD (G) TrkA bands normalized against DMSO control represented as averages ± SD (N=3).

### N-cadherin and TrkA show distinct expression in trigeminal neurons

To investigate possible interactions between N-cadherin and TrkA in chick trigeminal sensory neurons, we next examined the expression pattern of TrkA, and whether it colocalized with N-cadherin, both in the embryo and in trigeminal ganglion explant cultures. Immunostaining was used to evaluate spatial distribution within trigeminal neurons, and immunoblotting was conducted to assess protein levels and sizes. In the embryo, all trigeminal neurons express N-cadherin (Fig. 3A, A’), while only a subset express TrkA (Fig. 3B, B’), as expected (Halmi et al., 2025; Leonard et al., 2022). We noticed colocalization of the two proteins in certain areas within trigeminal neurons *in vivo*, including at the cell membrane and in axons (Fig 3C, C’, arrowheads). In trigeminal ganglia explant cultures, we saw that the majority of N-cadherin staining was found at the membrane (Fig. 3D, G), while the majority of TrkA staining was cytoplasmic (Fig. 3E, H). However, we also observed certain areas within these explants where N-cadherin and TrkA appeared to colocalize at the plasma membrane and intracellularly in apparent perinuclear vesicles (Fig. 3F, I, arrowheads).

**Figure 3:**
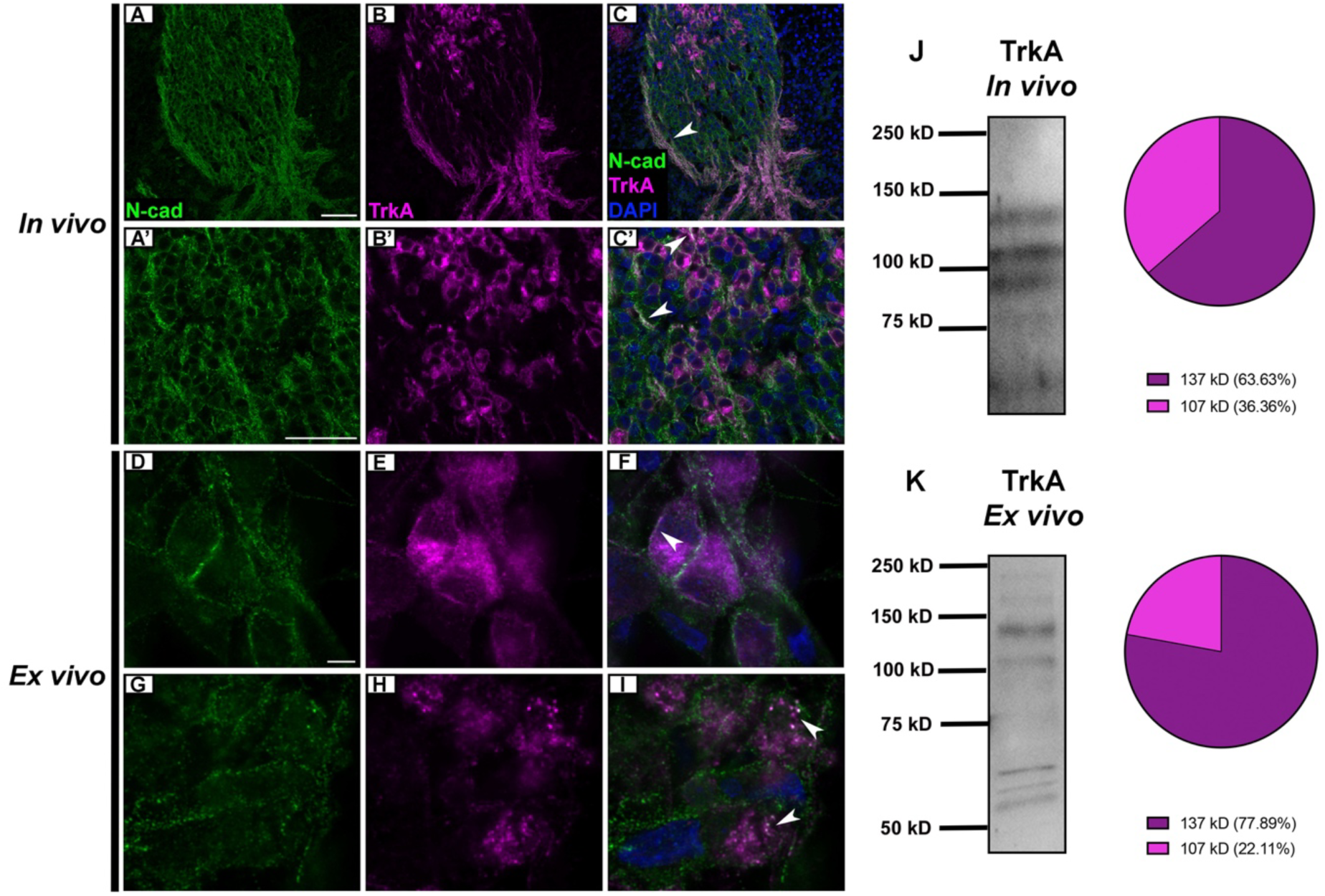
N-cadherin and TrkA are coexpressed in trigeminal neurons. (A-C’) Representative transverse sections through the trigeminal ganglion of an E5.5/HH28 chick embryo following immunohistochemistry for N-cadherin (N-cad) and TrkA. Scale bar in (A) is 50 µm and applies to (B, C). Scale bar in (A’) is 50 µm and applies to (B’, C’). (D-I) Representative trigeminal ganglia explant cultures after immunocytochemistry for N-cadherin and TrkA. Scale bar in (D) is 25 µm and applies to (E-I). Arrowheads indicate colocalization of N-cadherin and TrkA. (J, K) Immunoblots of HH28-30 trigeminal ganglia for TrkA with lysate from the embryo (*in vivo*) (J) and from trigeminal ganglia explant cultures (*ex vivo*) (K). Pie charts represent percentage of total signal coming from the different bands of TrkA.

When examining TrkA by immunoblotting, we observed similar banding patterns and molecular weights *in vivo* (Fig. 3J) and *ex vivo* (Fig. 3K) - 76 kD, 93 kD, 107 kD, and 137 kD - as observed previously (Fig. 1). We also noted two higher molecular weight bands at 223 kD and 177 kD in our *ex vivo* lysate, as seen in our glycan enrichment experiments (Fig. 1B). Collectively, these findings indicate that TrkA colocalizes with N-cadherin in trigeminal neurons and maintains similar, but not identical, banding patterns both *in vivo* and *ex vivo*.

### N-cadherin biochemically interacts with TrkA in trigeminal neurons

Due to the colocalization of N-cadherin and TrkA that we observed by immunostaining, we sought to determine whether these two proteins physically interact in the chick trigeminal ganglion. To test this, we used a co-immunoprecipitation assay and examined neuronal cell bodies separately from their axonal projections, as we hypothesized that different forms of TrkA would be expressed in neuronal cell bodies where protein processing occurs versus axons where the mature receptor functions in signaling with NGF. Trigeminal ganglia and their nerve branches were dissected from chick embryos, and once removed, a crude cut was made at the base of the OpV and MmV lobes to separate trigeminal cell bodies from axons (Fig. 4A, dotted lines). Once lysates were generated for trigeminal cell bodies and axons, an N-cadherin immunoprecipitation with a validated N-cadherin antibody (Halmi et al., 2022) was performed followed by TrkA immunoblotting to detect possible interactions. In the cell bodies experiment, we observed the 223 kD, 137 kD, and 107 kD TrkA bands in the input, all of which interact with N-cadherin (Fig. 4B). This differed subtly from what we noted in the axons, where only the 223 kD and 137 kD bands were present in the input and after N-cadherin immunoprecipitation (Fig. 4C). Interestingly, in both immunoprecipitations, we also observed two smaller TrkA bands at 75 kD and 58 kD. These results align with our hypothesis and indicate that N-cadherin interacts with different forms of TrkA within different compartments of trigeminal neurons.

**Figure 4:**
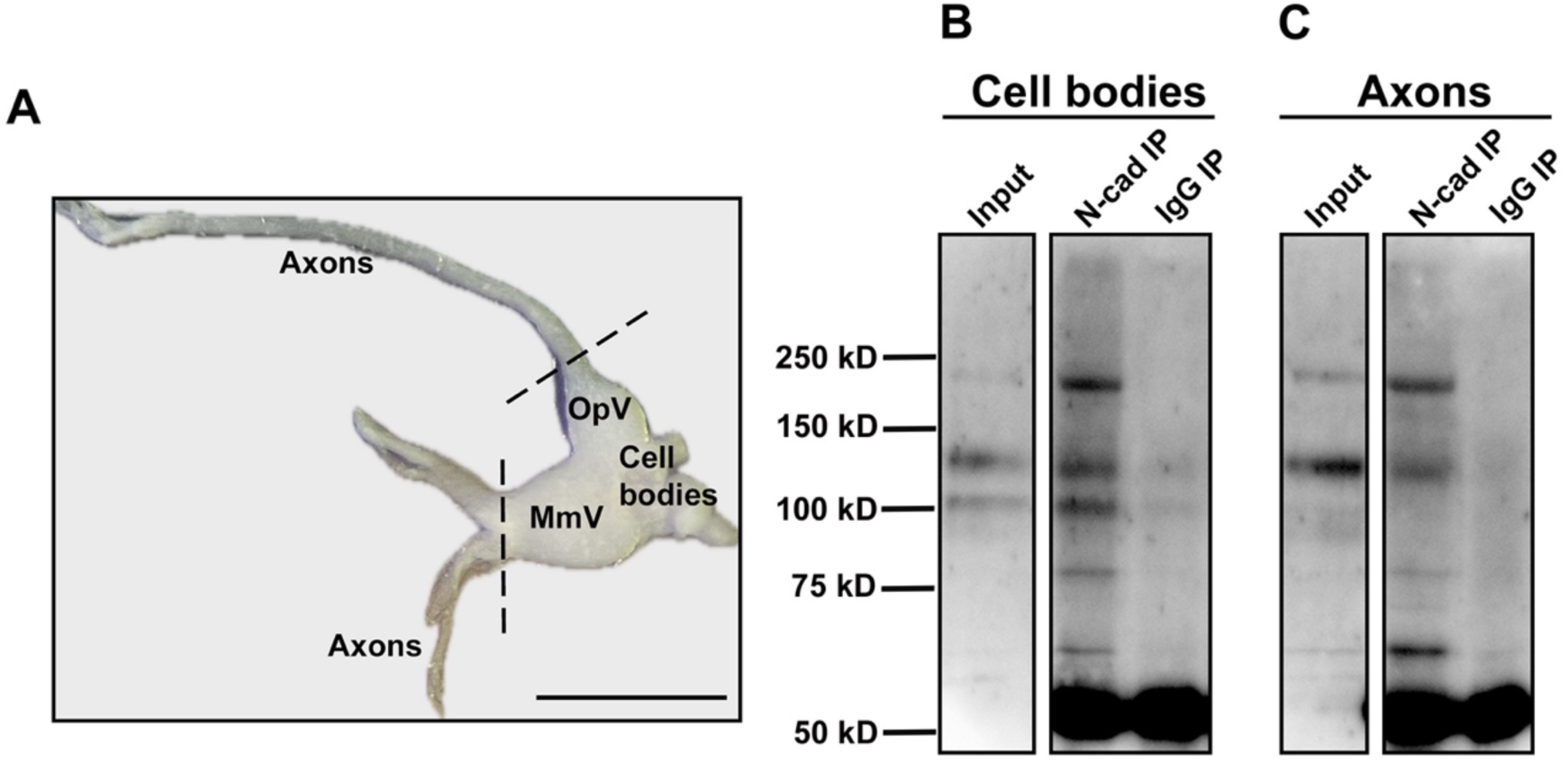
Different glycosylated forms of TrkA physically interact with N-cadherin in distinct compartments of the trigeminal ganglion. (A) Dissected trigeminal ganglion from an E5.5/HH28 chick embryo. Dotted lines indicate approximate location where cuts were made to separate trigeminal cell bodies from axons. Scale bar is 1mm. (B-C) N-cadherin-TrkA co-immunoprecipitation on pooled HH28-30 trigeminal cell bodies and axons. Input lanes contain 10% of lysate loaded in immunoprecipitation lanes. OpV, ophthalmic; MmV, maxillomandibular; IP, immunoprecipitation; IgG, Immunoglobulin G. N=3.

### TrkA and N-cadherin interact on the membranes of trigeminal neurons

Our results demonstrate that N-cadherin interacts with the 107 kD N-linked glycosylated, 137 kD N-linked and sialylated, and 223 kD glycoforms of TrkA. However, because only the fully glycosylated form of TrkA is thought to function at the neuronal plasma membrane in signaling with NGF, these results suggest that the interactions we observed between N-cadherin and TrkA are occurring at least partially intracellularly, and potentially in the organelles in which these modifications take place. To further explore this, we separated membrane-bound and membrane-associated proteins from cytosolic proteins in E6.5/HH28-30 trigeminal ganglia. We first verified the accuracy of our extraction method by probing for different proteins and examining in which fraction they were observed. Both N-cadherin and TrkA appeared solely in the membrane-extracted fraction (Fig. 5A). As receptors, these proteins are primarily expressed on the plasma membrane but would also be expected to be embedded in membranes when moving through the secretory pathway. Next, we looked at Beta-tubulin III (Tubb3), a neuronal microtubule protein that provides cytoskeletal support. Tubb3 was primarily expressed in the cytosolic fraction, although there was a small amount also found in the membrane fraction, which is not unexpected since it interacts with membrane-associated proteins and is found in the mitochondrial membrane of certain cells (Parker et al., 2022; Radwitz et al., 2022). Lastly, we examined Golgi matrix protein 130 (GM130) and SERCA2, membrane proteins expressed in the Golgi and ER, respectively. Both markers were enriched in the membrane fraction (Fig. 5A). This experiment confirms that our membrane extraction of trigeminal ganglia isolated membrane-bound and membrane-associated proteins from both the plasma membrane and membranes of organelles.

**Figure 5:**
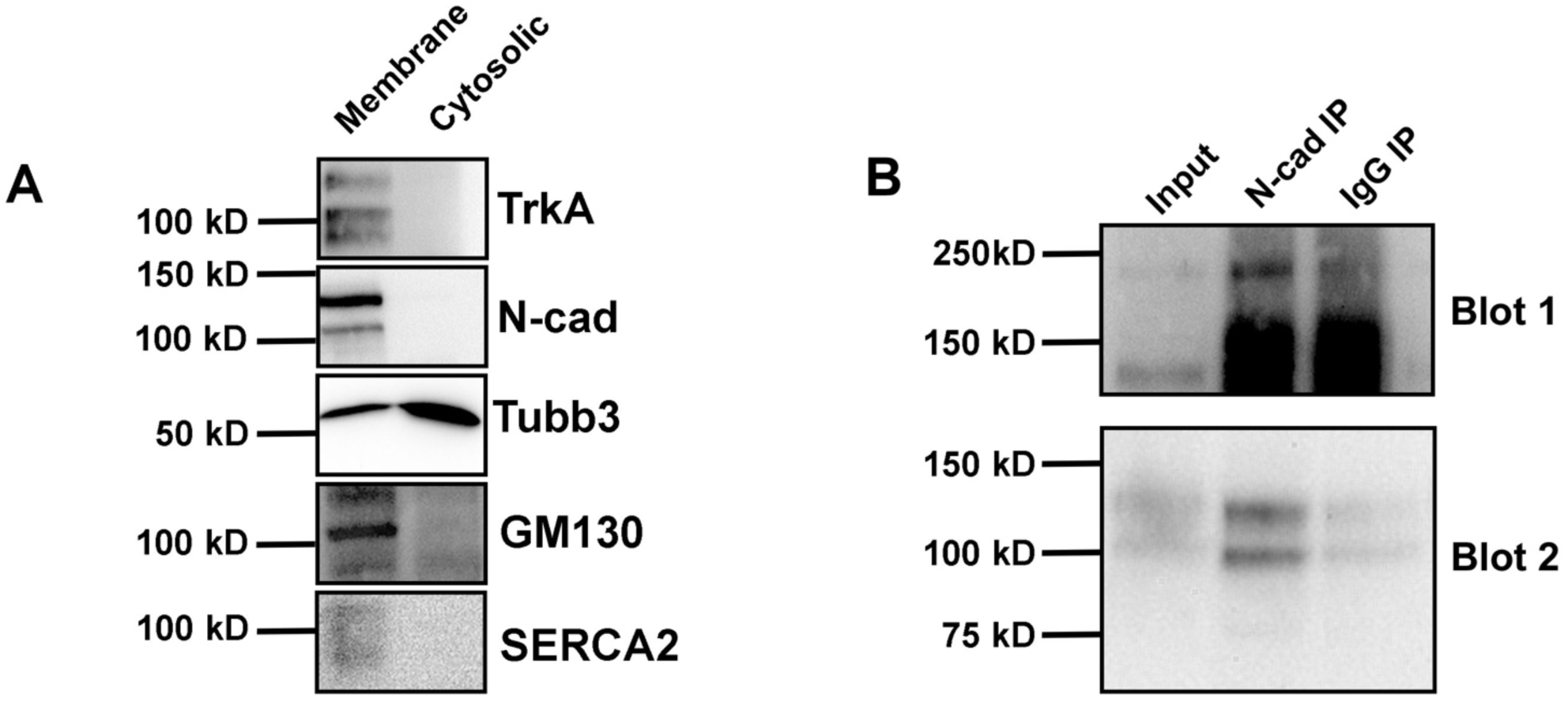
TrkA and N-cadherin interact on neuronal membranes. (A) Immunoblots of E6.5/HH28-30 trigeminal ganglia lysate following Mem-PER membrane extraction protocol for TrkA, N-cadherin, Tubb3, GM130 (Golgi protein), and SERCA2 (ER protein) using membrane-extracted and cytosolic-extracted protein fractions. (B) N-cadherin-TrkA co-immunoprecipitation on membrane-extracted fraction. Input contains 10% of lysate loaded in immunoprecipitation lanes. The top blot image is from one experiment using a high exposure and the bottom blot image is from a separate experiment with a lower exposure. IP, immunoprecipitation, IgG, Immunoglobulin G (N=3).

We next performed an N-cadherin immunoprecipitation on the membrane-extracted fraction followed by immunoblotting for TrkA. We observed an interaction between N-cadherin and the 107, 137, and 223 kD forms of TrkA (Fig. 5B). Given data on the partially glycosylated form of TrkA (107 kD in our experiments) from PC12 cells as well as the results from our Brefeldin A treatment (accumulation of the 107/98 kD bands after blocking ER to Golgi transport, Fig. 2), these results suggest that the 107 kD TrkA-N-cadherin interaction is occurring on intracellular membranes. Because sialic residues are added to TrkA in the Golgi, it is possible that the interaction between N-cadherin and the 137 kD band of TrkA is occurring in the Golgi, at the plasma membrane, or in both locations. This experiment demonstrates that the interactions between N-cadherin and TrkA are occurring on neuronal membranes and are likely happening both intracellularly and at the plasma membrane.

## Discussion

Although post-translational modifications of TrkA have been thoroughly examined in cell lines, whether these same modifications occur *in vivo* remain poorly understood. In this study, we explored the glycosylation of TrkA in chick trigeminal neurons. Our experiments led us to confirm that the 107 kD and 137 kD bands commonly associated with TrkA in the literature are an N-linked partially glycosylated form and a N-linked and sialylated fully glycosylated form, respectively. Through our experiments, we also observed multiple lower molecular weight forms of TrkA, as well as changes to TrkA after probing for O-linked glycans. Using this characterization, we next examined interactions between TrkA and the cell adhesion molecule N-cadherin and found that N-cadherin interacts with both partially and fully glycosylated versions of TrkA, as well as a higher and two lower molecular weight forms of TrkA. These interactions were observed in both trigeminal neuron cell bodies and axons, and on neuronal plasma membranes and intracellular membranes, highlighting a previously uncharacterized interaction between neurotrophic signaling receptors and cell adhesion that underlies trigeminal ganglion neurodevelopment.

### Maturation of chick TrkA involves multiple glycan modifications

To evaluate the identity of the TrkA bands observed in chick trigeminal ganglia, we first performed a series of experiments centered around perturbing or enriching for glycan residues on TrkA. Treatment of trigeminal ganglia lysate with PNGase F, which cleaves the glycosidic bond in N-linked glycoproteins, yielded one strong 88 kD band, confirming that the 93 kD, 107 kD, and 137 kD TrkA bands observed in the input lane are all N-linked glycosylated. In addition, we observed a 67 kD and a 56 kD band, likely the 76 kD and 59 kD TrkA bands observed prior to treatment, respectively. Whether these additional bands are intermediate forms of TrkA as it undergoes glycosylation or cleaved fragments of TrkA is not known. TrkA, along with other RTKs, is proteolytically processed to generate a soluble ectodomain fragment that is approximately 100 kD and a membrane-bound fragment of 41 kD (Cabrera et al., 1996; Díaz-Rodríguez et al., 1999). Based on this, it is possible that the 93 kD TrkA we observed is the soluble ectodomain of TrkA; however, caveats exist with this interpretation. After cleavage, the shed ectodomain diffuses away from the plasma membrane. For our experiments to detect this TrkA fragment, it would need to remain closely associated with the trigeminal ganglion. This is plausible, though, given how compact the neurons are at this stage in development and the potential for “trapping” the TrkA shed ectodomain in the extracellular matrix and mesenchyme surrounding the ganglion. Although intracellular cleavage fragments of TrkA remain poorly characterized, evidence of such processing has been reported in the human receptor (Merilahti et al., 2017; Merilahti & Elenius, 2019). However, we do not believe any of the TrkA bands observed in our experiments are intracellular fragments since our TrkA antibody recognizes an epitope in the extracellular domain.

Interestingly, the 53 kD band was the only form of TrkA that did not change after PNGase F treatment, suggesting that it lacked N-linked glycans. Neuraminidase treatment altered only the 137 kD and 93 kD bands, both of which endured a slight reduction in molecular weight, indicating that these are the only sialylated forms of TrkA. This result was surprising, because if sialylation were the only additional modification occurring to transform 107 kD TrkA to the 137 kD form, a total collapse of the 137 kD band into the 107 kD band after neuraminidase treatment would be expected. However, this was not the case, suggesting other modifications, such as O-linked glycosylation, could be occurring in addition to sialylation to generate this 137 kDa form. Alternatively, it is possible that the neuraminidase enzyme might not have access to all the sialic acid moieties in TrkA due to the complexity of the protein structure, or our treatment time was insufficient to remove every sialic acid. Even with these considerations, our results mirror those observed for TrkA after neuraminidase treatment in PC12 cells, where the 137 kD band also was not completely reduced to the 107 kD band (Jullien et al., 2002).

We next sought to test whether O-linked glycosylation was present on TrkA. While the TrkA bands observed after O-glycosidase + neuraminidase treatment were almost identical to neuraminidase treatment alone, surprisingly, a very prominent 177 kD band was also evident after O-glycosidase + neuraminidase treatment. This band likely arises from enzymatic alterations to the 223 kD form of TrkA that we sometimes observe in the input control (Supplemental Fig. 1) as well as in the N-cadherin-TrkA co-immunoprecipitation. Based on the size of this band in combination with what is reported in the literature (Hartman et al., 1992), it is possible that this is a NGF-TrkA complex containing Core 1 and 3 O-glycans. The PNGase F + neuraminidase treatment yielded the same result as the PNGase F only treatment, with one pronounced band at 88 kD along with lower molecular weight bands at 67 kD, 56 kD, and 53 kD. This result indicates that TrkA is heavily N-linked glycosylated, and N-glycans serve as the base upon which terminal glycans, including sialic acids, are built. While the treatment with all three enzymes yielded bands observed in other treatments, interestingly, we detected a 52 kD band that was not found in any other condition. One possible explanation is that the combined enzymatic treatment enhanced glycan accessibility, allowing the enzymes to act with greater efficiency. This form of TrkA might be the nascent protein, or a very slightly modified form of TrkA shortly after synthesis. Given the dynamics of protein translation, it is possible that we were unable to capture the truly unmodified form of TrkA as often these modifications are added as proteins are being translated (Helenius & Aebi, 2001). Furthermore, our incubations may not have given the enzymes sufficient time to cleave all the residues from TrkA. However, we do not think this is the case since previous work performed PNGase F treatment for up to 36 hours and still did not uncover the “core” TrkA protein (Watson et al., 1999). It is equally plausible that other types of post-translational modifications are present on these TrkA forms that are unrelated to glycosylation. In addition to the importance of phosphorylation on TrkA function, ubiquitination regulates TrkA and therefore could be contributing to these changes in molecular weight (Georgieva et al., 2011; Yu et al., 2014). Interestingly, our input control and mock control samples yielded slightly different results, with the 223 kD form of TrkA only present in the former. This could be due to the presence of additional reducing agents (SDS and DTT) in the latter’s denaturation buffer, which, along with the Laemmli buffer, cause this higher molecular weight complex to come apart.

To further parse out these different glycans, we performed lectin binding assays. While ConA binds to alpha-linked mannose residues, WGA recognizes N-acetyl-glucosamine groups and sialic acids. Therefore, we hypothesized that ConA and WGA enrichment would enhance the 107 kD, N-linked glycosylated and 137 kD sialylated TrkA bands, respectively. Interestingly, both bands were enriched with both treatments. Although this is in contrast to similar experiments performed in PC12 cells (Jullien et al., 2002), this difference can be explained by the lysate source. The embryonic environment is more complex and heterogenous than cell lines; therefore, it is possible that TrkA glycosylation within the embryo contains additional and/or different glycans. Next, we used PNA, which binds Galactosyl-β(1-3)-N-acetyl-galactosamine, a type of O-linked glycan, based on other work that found O-linked glycans on TrkA (Lin et al., 2022). Surprisingly, a 117 kD TrkA band was enriched after PNA treatment. Because this band was not observed in these other treatments and did not appear in our enzyme assays, we believe that this is an intermediate version of TrkA that is expressed at low levels. Surprisingly, ConA, WGA, and PNA all enriched the 177 kD band observed after O-glycosidase treatment. Thus, this band contains both N- and O-linked glycans. Although this 177 kD band also appeared when all three enzymes (PNGase F, neuraminidase, and O-glycosidase) were combined, it is possible that the enzymatic digestion time was insufficient, or that this form of TrkA contains complex secondary structures that make it difficult for PNGase F and neuraminidase to properly access all moieties. Another explanation is that the 177 kD form of TrkA contains Core 2 and Core 4 O-glycans, which do not get cut by O-glycosidase. A more thorough analysis of these TrkA glycoforms through mass spectrometry would be beneficial to further classify the profile of these bands.

In concert with these experiments, we treated trigeminal neuronal cultures with inhibitors of the secretory pathway to define how these perturbations impacted the processing of TrkA. Although the molecular weights observed for TrkA in this experiment (98 kD and 122 kD) were slightly different that those observed *in vivo* and our other *ex vivo* experiments (107 kD and 137 kD), we believe they represent the same TrkA glycoforms and are just disparities due to the nature of SDS-PAGE. Monensin inhibits transport out of the Golgi by disrupting ion gradients. After immunostaining, we noted a near total accumulation of TrkA in a perinuclear location, which we expect is the Golgi. While little is known about TrkA sialylation, sialic acids are added to proteins as they move through the Golgi stacks (Zhu et al., 2024). Therefore, we would not expect the fully glycosylated 137 kD form of TrkA to be observed in the Monensin-treated group, but rather only the 107 kD form that is accumulating within the Golgi apparatus. Brefeldin A disrupts anterograde transport from the ER to the Golgi and causes disruption of the Golgi complex (Reaves & Banting, 1992). This treatment led to diffuse TrkA staining throughout the soma, and a robust 107 kD band after immunoblotting, suggesting that, in chick trigeminal neurons, the 107 kD partially glycosylated form of TrkA is in the ER, and further confirming that sialylation occurs largely within the Golgi. The diffuse staining of TrkA likely corresponds to the partially glycosylated form being dispersed throughout the ER of the cell, and similar results have been observed after Brefeldin A treatment in other types of neurons (Jareb & Banker, 1997). Lastly, Tunicamycin treatment, which prevents all N-linked glycosylation from occurring, led to accumulation of TrkA throughout the soma cytoplasm, and a near complete loss of both the 107 kD and 137 kD TrkA bands. Tunicamycin induces ER stress, protein misfolding, and eventual cell death (Heifetz et al., 1979). In keeping with this, TrkA immunostaining appears dim and punctate, suggesting unhealthy cells and protein processing. While the 107 kD and 137 kD bands were diminished after Tunicamycin treatment, very interestingly, that treatment group was the only condition in which a lower molecular weight band of 80 kD accumulates, supporting the notion that this could be an “immature” form of TrkA that possesses little to no glycosylation. These findings could reflect the differences noted for TrkA immunoblotting banding patterns *in vivo* and *ex vivo* and/or indicate that the Tunicamycin treatment time was not sufficient to remove all N-glycans and reveal the true core protein, especially since the 137 and 107 kD bands are still present. One main limitation of this experiment is the lack of available organelle markers compatible with immunostaining of chick tissue. Having clear markers to label organelles would be an excellent addition to these studies to verify the subcellular location of TrkA accumulation within the cells.

### TrkA interacts with N-cadherin dynamically in different compartments of trigeminal neurons

We next sought to examine the coexpression of TrkA with N-cadherin in E5.5/HH28 chick embryos and found that both *in vivo* and *ex vivo*, the two colocalized along the plasma membrane of trigeminal neurons. Although it was difficult to detect intracellular colocalization via immunostaining in our *in vivo* samples due to the compact nature of the trigeminal ganglion at this stage of development, we did observe intracellular colocalization of TrkA and N-cadherin *ex vivo,* likely within vesicles and in perinuclear structures. Given that we observed TrkA and N-cadherin colocalization in both the cell bodies and along the axons, we chose to explore whether these two proteins were interacting in both compartments. Through co-immunoprecipitation assays, we found that N-cadherin interacts with the 107 kD and 137 kD form of TrkA in cell bodies but only the 137 kD form of TrkA in axons. TrkA synthesis occurs in the cell bodies of neurons as it progresses through the secretory pathway to transform from an immature protein into a fully glycosylated receptor. Only then does it undergo anterograde trafficking through the axon to be inserted into the axonal plasma membrane, where it acts as a signaling receptor for NGF. Thus, the absence of the 107 kD TrkA band from both the input and IP lanes of our axon sample is not surprising and suggests that this band is a partially glycosylated form of TrkA moving through the secretory pathway within the cell bodies. Because mature TrkA interacts with NGF on the axon terminals, an interaction between the 137 kD mature form of TrkA and N-cadherin in the cell bodies is likely occurring on the plasma membrane or within vesicles either traveling anterogradely or retrogradely from axons. Experiments in rodent sympathetic ganglia have shown that transcytosis can cause the mature TrkA receptor to be endocytosed from the surface of cell bodies and anterogradely trafficked to the axon surface (Ascaño et al., 2009; Connor et al., 2023; Yamashita et al., 2017). It is possible that this process is also occurring in chick sensory ganglia. In both compartment immunoprecipitations, we observed a larger molecular weight band at 223 kD in addition to two smaller 76 kD and 59 kD bands. Previous studies noted a band size of 220 kD in PC12 cells that likely represents a NGF-TrkA protein complex based on radiolabeling (Hartman et al., 1992). As such, the 223 kD band in our experiments could represent an NGF-bound TrkA-N-cadherin complex, while the lower molecular weight bands could be differently glycosylated forms of TrkA interacting with N-cadherin as they move through the secretory pathway.

While our co-immunoprecipitation experiments confirm an interaction between N-cadherin and multiple different forms of TrkA, whether they are directly interacting extracellularly, intracellularly, or indirectly via a complex remains unanswered. Cadherin-RTK interactions contribute to cellular processes underlying both development and disease, which have given insight into how these receptors regulate each other. Typically, cadherins and RTKs directly interact via their extracellular domains, as noted in E-cadherin-EGFR and N-cadherin-FGFR interactions (Andl & Rustgi, 2005; T. Nguyen & Mège, 2016; Qian et al., 2004). Moreover, in some cases, the binding of the cadherin to the RTK limits the ability of ligand binding (Qian et al., 2004; Blaschuk, 2025), although we do not believe this to be the case in our experiments given the presence of the 223 kD TrkA band that is purported to possess NGF. Conversely, N-cadherin is also capable of preventing FGFR from getting internalized after ligand binding, keeping it constitutively active at the membrane to promote metastasis (Suyama et al., 2002). Another possibility is that N-cadherin and TrkA are not interacting directly with each other and are instead forming a complex through adaptor proteins. Two potential candidates for this are Shc and ß-catenin, as demonstrated in cell lines. Shc, a common adaptor protein for neurotrophic signaling, binds the C-terminus of N-cadherin (Xu et al., 1997) while ß-catenin, a crucial connector of cadherins to the actin cytoskeleton, interacts with Trk receptors (David et al., 2008). Future studies would aid in addressing whether the interaction between N-cadherin and TrkA is direct or indirect.

### TrkA and N-cadherin interactions occur both at the cell membrane and intracellularly

We lastly sought to explore where the interactions between N-cadherin and TrkA seen from Figure 4 were occurring within the trigeminal neurons. To test this, we separated all membrane-bound and membrane-associated proteins from cytosolic proteins and performed an immunoprecipitation for N-cadherin followed by immunoblotting for TrkA. Both the partially and fully glycosylated forms of TrkA were found to interact with N-cadherin in the membrane fraction, as well as the 223 kD band which we hypothesize to be a NGF-TrkA complex. Given the cumulative knowledge surrounding cadherin-RTK interactions, and how they commonly regulate each other’s activity at the plasma membrane, it is not surprising to see the functional form of TrkA interacting with N-cadherin, either on the plasma membrane or within other membrane-possessing structures like vesicles (e.g., signaling endosomes) (Harrington & Ginty, 2013; Howe et al., 2001; Sharma et al., 2010). Furthermore, if the 223 kD band we observe is truly a NGF-TrkA complex that associates with N-cadherin, this interaction would likely be found at the plasma membrane or on the membrane of an internalized vesicle. Strikingly, we observed that the 107 kD, partially glycosylated form of TrkA also interacts with N-cadherin. Because only the 137 kD form of TrkA reaches the plasma membrane, this interaction with the 107 kD form likely occurs intracellularly, either on a membrane of the ER, Golgi, or within a transport vesicle. We can also not rule out that the interaction between the 137 kD form of TrkA and N-cadherin solely occurs on the plasma membrane. This interaction could also be taking place on the Golgi membrane after TrkA has been fully modified, or on a vesicle traveling to the plasma membrane. While there is very limited knowledge surrounding intracellular cadherin-RTK interactions, ER to Golgi transport of N-cadherin-FGFR complexes has been demonstrated (Nakamura et al., 2008). Thus, a similar phenomenon could be occurring between N-cadherin and TrkA as they travel through the secretory pathway. Future experiments examining the subcellular location of this interaction would give more insight into its functional role.

## Conclusion

In this study, we identified different glycoforms of the neurotrophic receptor, TrkA, in chick trigeminal sensory neurons and demonstrate that these forms of TrkA interact with N-cadherin on the membranes of neurons. These results contribute to our general understanding of how post-translational modifications and membrane interactions regulate TrkA localization during sensory development. Given the integral roles that both TrkA and N-cadherin play during neuronal development, their interaction likely contributes to the coordinated assembly of the trigeminal ganglion. As aberrant RTK and cadherin signaling are increasingly becoming recognized in pathological conditions, and subsequently contribute to altered cell fate and proliferation, understanding how these pathways intersect during normal development can provide important knowledge on how their dysregulation contributes to diseases. Future studies assessing how N-cadherin modulates TrkA trafficking, signaling, and localization at the membrane will clarify the mechanisms behind adhesion-dependent signaling that shapes the sensory nervous system.

## Supplemental Figures

**Supplemental Figure 1:**
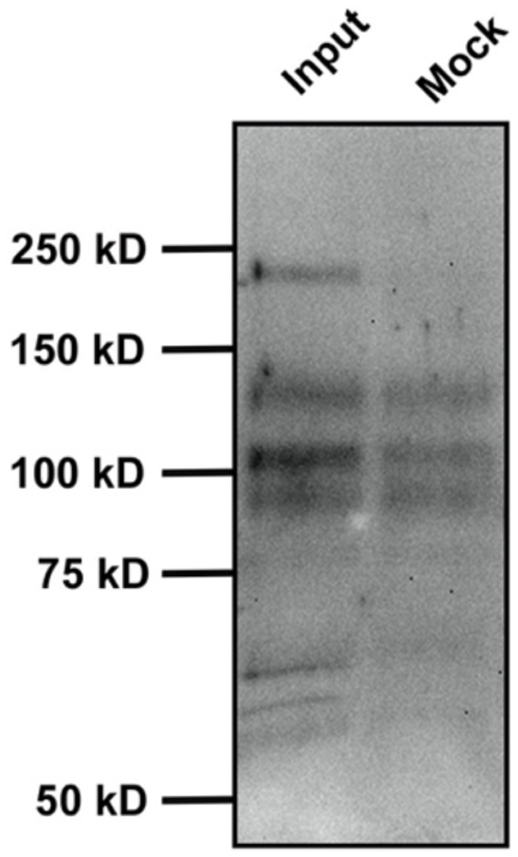
Higher molecular weight TrkA is observed in trigeminal ganglia lysate. E6.5/HH28-30 trigeminal ganglia lysate treated with enzymes to inhibit glycosylation followed by immunoblotting for TrkA. Input: Untreated trigeminal ganglia lysate. Mock (control): Lysate mixed with enzyme buffers (N=3).

**Supplemental Figure 2:**
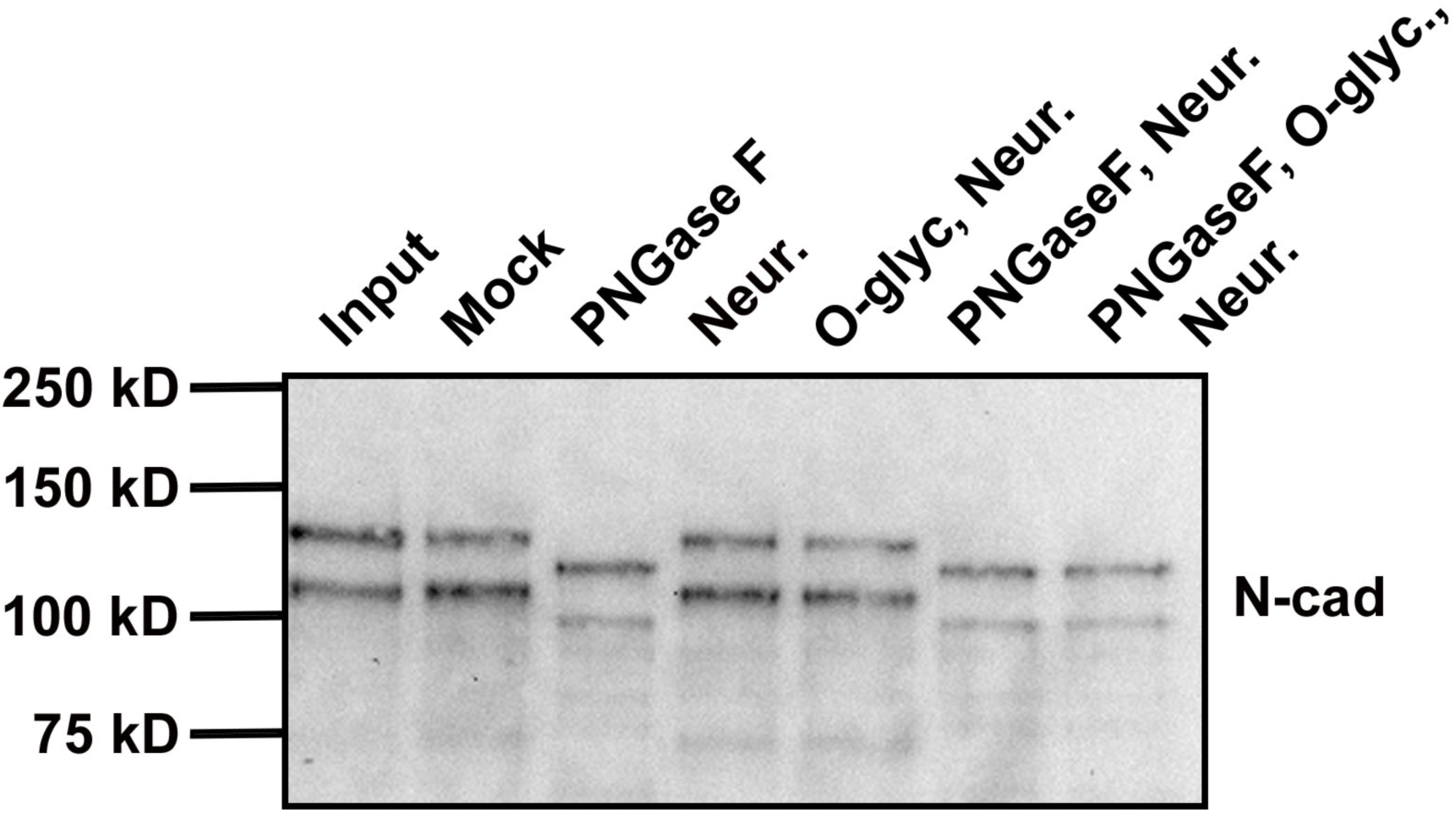
N-cadherin is N-linked glycosylated. E6.5/HH28-30 trigeminal ganglia lysate treated with enzymes to inhibit glycosylation followed by immunoblotting for N-cadherin (N-cad). Input: Untreated trigeminal ganglia lysate. Mock (control): Lysate mixed with enzyme buffers. PNGase F, Neuraminidase, O-glycosidase (O-glyc.) (N=2).

